# Mechano-chemical feedback leads to competition for BMP signalling during pattern formation

**DOI:** 10.1101/2020.09.23.309435

**Authors:** Daniel J. Toddie-Moore, Martti P. Montanari, Ngan Vi Tran, Evgeniy M. Brik, Hanna Antson, Isaac Salazar-Ciudad, Osamu Shimmi

**Affiliations:** Institute of Biotechnology, University of Helsinki, 00014 Helsinki, Finland; Institute of Molecular and Cell Biology, University of Tartu, 51010 Tartu, Estonia; Genomics, Bioinformatics and Evolution. Departament de Genètica i Microbiologia, Universitat Autònoma de Barcelona, 08193, Cerdanyola del Vallès, Spain; Centre de Rercerca Matemàtica, 08193, Cerdanyola del Vallès, Spain

## Abstract

Developmental patterning is thought to be regulated by conserved signalling pathways. Initial patterns are often broad before refining to only those cells that commit to a particular fate. However, the mechanisms by which pattern refinement takes place remain to be addressed. Using the posterior crossvein (PCV) of the *Drosophila* pupal wing as a model, into which bone morphogenetic protein (BMP) ligand is extracellularly transported to instruct vein patterning, we investigate how pattern refinement is regulated. We found that BMP signalling induces apical enrichment of Myosin II in developing crossvein cells to regulate apical constriction. Live imaging of cellular behaviour indicates that changes in cell shape are dynamic and transient, only being maintained in those cells that retain vein fate after refinement. Disrupting cell shape changes throughout the PCV inhibits pattern refinement. In contrast, disrupting cell shape in only a subset of vein cells can result in a loss of BMP signalling. In addition, we observed that expressing the constitutively active form of the BMP type I receptor in clones caused apical constriction autonomously and often induced BMP signalling loss in the PCV region in a non-autonomous manner. We propose that the cell shape changes of future PCV cells allow them to compete more efficiently for the basally localised BMP signal by forming a mechano-chemical feedback loop. This study highlights a new form of competition among the cells: competing for a signal that induces cell fate.

## Introduction

Pattern formation is a fundamental process in animal development, for which various molecular mechanisms have been proposed, including gene regulatory networks and growth factor signalling^1,2^. Developmental patterning often involves refinement from a broad initial area of competency for a fate to only those cells that commit to it, with neighbours losing competence and following an alternate fate path (Fig. 1a)^1,2^. Whilst some mechanisms of pattern refinement, such as transcriptional networks and lateral inhibition, have previously been investigated, the role played by diffusible growth factor signalling, in particular the interactions between signalling and morphogenesis, has been less explored^3,4^.

**Fig. 1:**
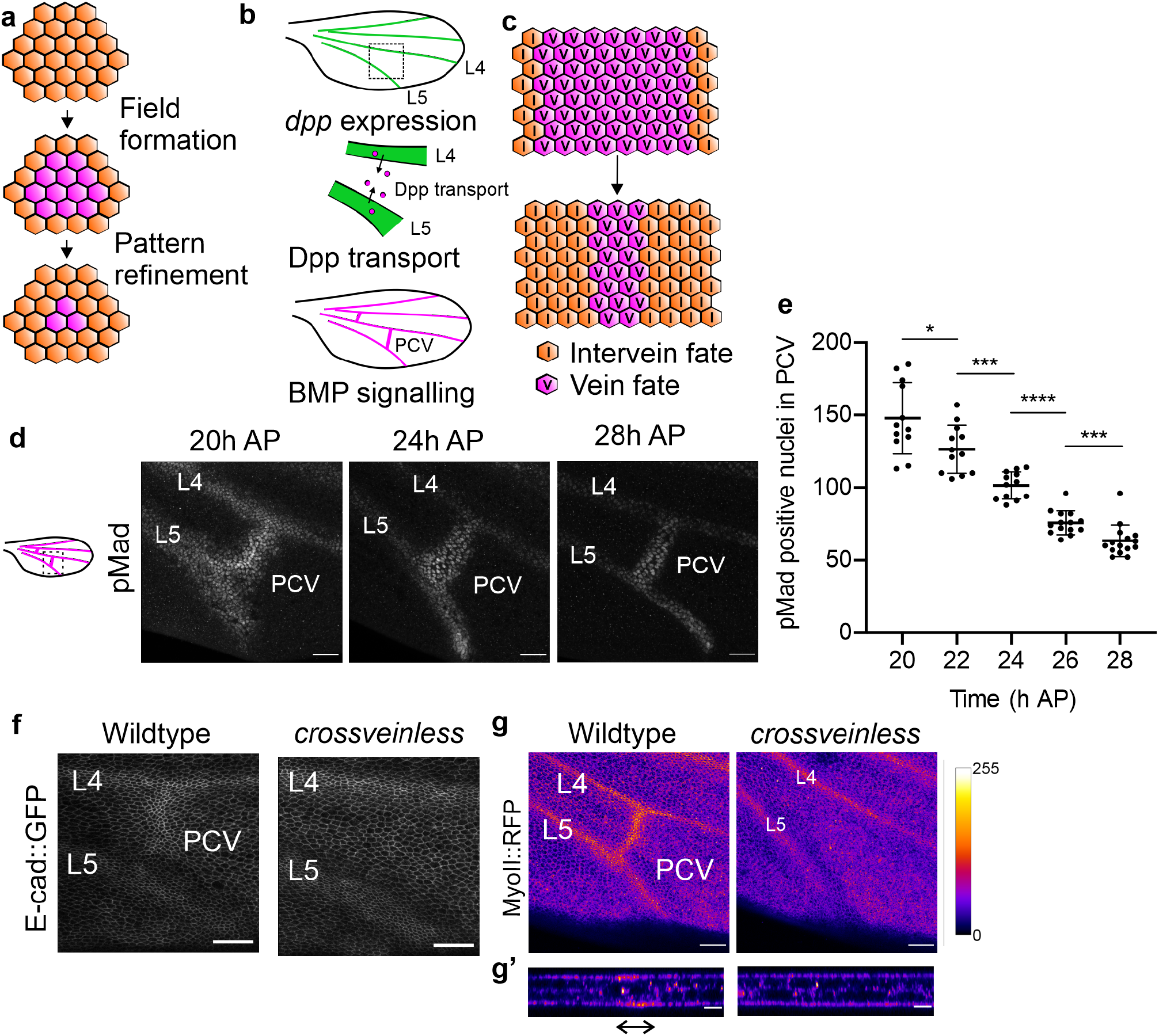
The PCV field refines during vein patterning and morphogenesis. **a,** Schematic depicting the refinement of developmental patterns from an initial field to cells committing to a fate. **b,** The expression pattern and signalling pattern of the BMP ligand Dpp in the developing pupal wing. L4, L5, and PCV denote longitudinal veins 4, 5, and posterior crossvein, respectively. Top: *dpp* mRNA (green) is expressed in longitudinal veins, but not in crossveins during early pupal stages. Middle: Schematic model of Dpp/BMP ligand transport from the longitudinal veins into PCV. Bottom: BMP signalling (magenta) is detected at all wing vein primordia including longitudinal veins and crossveins. **c,** Schematic depicting the refinement of the BMP signalling pattern in the PCV field. **d,** BMP signalling (shown by pMad) in the PCV field of wild type pupal wings at 20 h, 24 h and 28 h AP. Left: Schematic of pupal wing. Approximate position of imaging is shown as a square. Median filter applied. **e,** The number of cells in which BMP signalling is occurring during the refinement period. Sample sizes are 12 (20 h), 12 (22 h), 12 (24 h), 14 (26 h) and 15 (28 h). \**P* = 0.0252, 22-24h: \*\*\**P* = 0.0005, \*\*\*\**P* < 0.0001, 26-28h: \*\*\**P* = 0.0002. Data are mean + s.d. and were analysed by two-sided Mann-Whitney test. **f,** E-cad::GFP in the PCV region in wild type (left) and *crossveinless* mutant (right) pupal wings at 24 h AP. Apical cell shapes are highlighted by max composite of E-cad:GFP. **g,** Heatmap of the apical intensity of MyoII::RFP in cells of the PCV region in wildtype (left) and *crossveinless* mutant (right) pupal wings at 24h AP. Optical cross sections on the PCV region are shown at the lower panel (**g’**). Prospective PCV position is indicated by double-headed arrow. The apical distribution of MyoII is shown through the PCV region. Scale bars: 25 μm for **d**, **f** and **g,** and 10 μm for **g’**.

The posterior crossvein (PCV) of the *Drosophila* pupal wing serves as an excellent model to address the dynamics of signalling and morphogenesis, as its formation is initially directed by a single signalling pathway: Bone morphogenetic protein (BMP) signalling^5,6^. The *Drosophila* BMP ligand Decapentaplegic (Dpp) is initially expressed in the adjacent longitudinal veins (LVs) and is extracellularly transported into the prospective PCV region along the basal surfaces of the two cell layers that comprise the wing epithelia (Fig. 1b)^7^. Extracellular transport at this stage requires the BMP binding proteins Short gastrulation (Sog) and Crossveinless (Cv) which bind to the Dpp ligand and facilitate its active transport into the PCV region^8,9^.

BMP signalling induced by the Dpp ligand forms the PCV field, directing cells to become competent for vein, rather than the intervein fate that occurs if BMP signalling is not activated or is not sustained (Fig. 1c)^5,8^. Continuous extracellular Dpp transport seems to be crucial for a period of around 10 hours (18 – 28 h after pupariation (AP)) to maintain the PCV field and vein fate competence, before PCV cells begin to express the ligand themselves^5^. Continued extracellular signalling and vein morphogenesis occur concurrently, as morphogenesis begins shortly after BMP signalling is activated^6^. Refinement of the BMP signalling pattern during this time window has previously been observed; however, how pattern refinement takes place has not been addressed^10^. In this study, to understand how refinement of the BMP signalling takes place during PCV fate determination, we utilised *Drosophila* genetics and *in vivo* live imaging. Our data reveal that the dynamics of cell shape direct refinement by forming a mechano-chemical feedback loop that drives competition for the vein-fate-determining BMP signal.

## Results

### BMP signalling induces cell shape changes during pattern refinement

First, we confirmed that refinement of the PCV field takes place during PCV morphogenesis. The PCV field is defined as the cells in which BMP signalling occurs, as indicated by staining with anti-phosphorylated Mad antibody (pMad)^11^, and which are therefore competent to assume a vein fate. Our data reveal that the number of cells within the PCV field reduces between 20h AP, shortly after the initiation of PCV patterning, and 28h AP, when PCV cells express *dpp* themselves (Fig. 1d, e). Thus we term the period from 20 to 28 hours AP the ‘refinement period’.

During this period, apical constriction of vein cells appears to be the hallmark of vein morphogenesis^10,12,13^. Since BMP signalling is thought to initiate PCV development^7^, we next asked whether BMP signalling directs the wing vein-like cell shape changes (hereafter referred to as “cell shape changes”) that occur during PCV morphogenesis. To answer this question, we captured images of apical cell shapes in the PCV region of *crossveinless* mutant wings, into which Dpp cannot be transported, and where BMP signalling is thus inactive. We observed that apical constriction does not occur where the PCV normally forms (Fig. 1f)^8^, indicating that BMP signalling is required for the cell shape changes that occur during PCV morphogenesis.

The activity of Myosin II (MyoII) has been proposed to be the driving force behind cell shape dynamics such as apical constriction^14^. To investigate whether BMP signalling directs cell shape change through MyoII activity, we analysed the spatial localisation of MyoII using MyoII regulatory light chain (MRLC) tagged with RFP in wild type and *crossveinless* pupal wings^15^. In wild type wings, MyoII is enriched in the apical compartment of PCV cells, with lower basal levels, but in contrast, neighbouring intervein cells have lower apical levels of MyoII than the PCV (Fig. 1g, Supplementary Fig. 1a). Conversely, in *crossveinless* pupal wings, apical MyoII enrichment is not observed in the PCV region, although apical enrichment of MyoII is still detected in LVs (Fig. 1g, Supplementary Fig. 1a). These findings suggest that BMP signalling facilitates the apical localisation of MyoII to promote apical constriction of PCV cells. This was further confirmed by the observation that ectopic expression of the constitutively active form of BMP type I receptor Thickveins (Tkv^QD^) in mosaic analysis with a repressible cell marker (MARCM) clones within the pupal wing induces apical enrichment of MyoII and apical constriction (Supplementary Fig. 1b)^10,16^.

### Time lapse imaging during PCV morphogenesis reveals reversible fate path

As wing vein morphogenesis directed by BMP signalling and refinement of the BMP signalling pattern occur concurrently, we hypothesised that these events could be mutually coordinated. To address this, we employed *in vivo* live imaging of pupal wings expressing GFP-tagged E-cadherin (E-cad::GFP) to observe cell shape changes during the refinement period^17^. We tracked the apical shapes of cells that are part of the PCV at the end of the refinement period, and thus retained vein fate competence, and compared them to the cells flanking this region (Fig. 2a-c, Supplementary Videos 1-3). Intriguingly, whilst the cells that will form the PCV constrict apically throughout the refinement period, several of the cells immediately flanking these constrict apically at early time points, but fail to maintain their vein-like morphology at later time points, eventually reverting to an intervein fate (Fig. 2a-d, Supplementary Videos 1, 2).

**Fig. 2:**
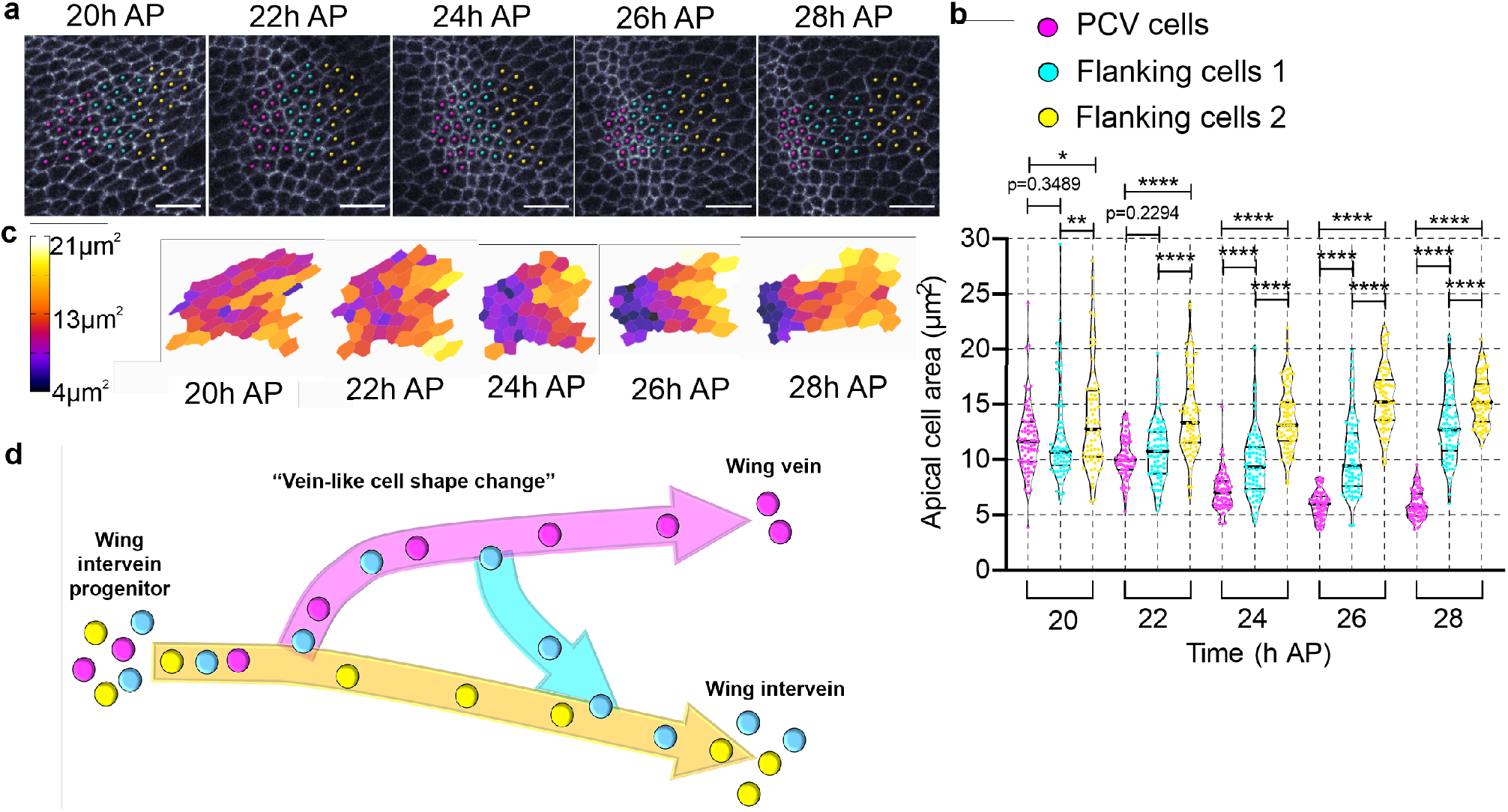
Changes in cell shape are dynamic and transient during pattern refinement. **a,** Time-lapse images of E-cad::GFP in the PCV region at 20 h, 22 h, 24 h, 26 h and 28 h AP. Three clusters of cells are marked. Future PCV cells (magenta) show progressive apical constriction. Cells immediately adjacent to future PCV cells (cyan) show transient apical constriction during 22 h and 26 h AP before reverting to an intervein-like structure. Cells further from future PCV cells (yellow) do not show apical constriction. PCV cells are categorised by their shape at 28 h AP and flanking categories by their relative position and shape at 28 h AP. Scale bars: 10 μm. **b,** Apical size of PCV cells and their neighbours during the refinement period. N=75 cells per category (15 cells tracked per category in each wing, for 5 wings). Violin plots show median, and 25th and 75th percentiles. Data from five independent time-lapse images. Each data point [PCV cells: magenta, cells adjacent to PCV (Flanking cells 1): cyan, cells further from PCV (Flanking cells 2): yellow] represents one cell. \**P* = 0.0185, \*\**P* = 0.0071, \*\*\*\**P* < 0.0001. Data were analysed by two-sided Mann-Whitney test. **c,** Heat map showing the changes in apical area of cells of the PCV field. The heatmap was produced using the ROI colour coder plugin, part of the BAR collection of ImageJ. The same cells are shown as in 2a. **d,** Schematic of changes in fate path during PCV patterning. Cells that are initially on a vein-like cell shape (cyan) lose competence during patterning, and move outside of the vein fate path (magenta) into the intervein fate path (yellow).

### Cell shape change and pattern refinement are coupled

We hypothesised that cell shape changes themselves may affect signalling pattern refinement and thus cell fate choice in the PCV region. To test this idea, we modulated cell shape changes in the developing wing by attenuating MyoII activity using a dominant negative form of the Myosin Heavy Chain (MyoII-DN)^18^. Inhibiting MyoII activity across the posterior wing blade for 10 hours during PCV morphogenesis is sufficient to disrupt apical constriction in the PCV region and LV cells of 23h AP pupal wings (Fig. 3a-c). Intriguingly, loss of MyoII activity throughout the posterior wing results in a broader range of BMP signalling in the PCV region than in control wings, suggesting that cell shape changes are necessary for the refinement of the BMP signalling, but not for BMP signalling itself (Fig. 3d, e). Consistently, adult wings show disrupted PCV patterning, manifesting both as vein thickenings and branches that extend from the normally straight PCV, suggesting that the organisation of cell fates has been disrupted (Fig. 3f). Additionally, BMP signalling was missing in the PCV region when MyoII activity was disrupted in the posterior half of *crossveinless* wings, indicating that the unrefined BMP signalling pattern is still being directed by extracellular BMP signalling (Supplementary Fig. 2a). We further confirmed that dynamic MyoII activity is crucial for refinement of BMP signalling using the ectopic expression of constitutively active Myosin binding subunit (Mbs), a regulatory subunit of myosin phosphatase, which also leads to a broader range of BMP signalling in the PCV region (Supplementary Fig. 2b, c)^19^.

**Fig. 3:**
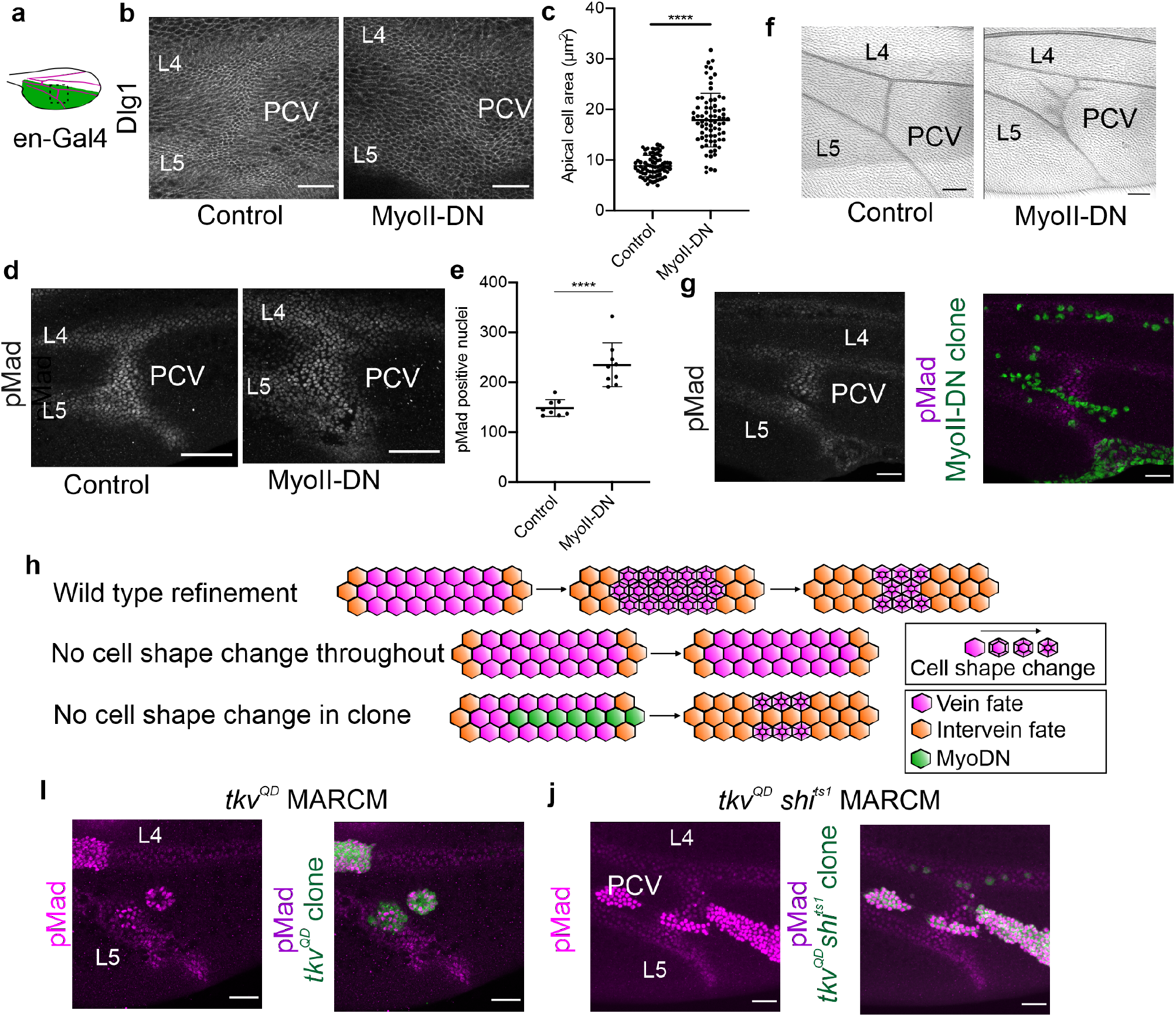
Loss of MyoII activity has context-specific effects on BMP signalling pattern refinement. **a,** Schematic of pupal wing. Posterior region of the wing and Engrailed-Gal4 expression pattern is shown in green. Approximate position of imaging is shown as a square (dotted outline). **b,** Apical cell shape (Dlg1) in the PCV region in control (left, *en > mCD8::GFP*) and MyoII attenuated pupal wings (right, *en > MyoII-DN*) at 23 h AP. **c,** Apical size of cells in which BMP signalling is occurring (and thus are within the PCV field) in the PCV of 23h AP pupal wings. N=75 cells per category (15 cells in each wing, for 5 wings). \*\*\*\**P* < 0.0001. Data are mean ± s.d. and were analysed by two-sided Mann-Whitney test. **d,** pMad expression in the PCV region in control (left, *en > mCD8::GFP*) and MyoII attenuated pupal wings (right, *en > MyoII-DN*) at 23 h AP. MyoII-DN was expressed throughout the posterior wing by *en-Gal4* for 10 hours prior to dissection. Median filter applied. **e,** Number of cells in which BMP signalling is occurring (and thus are within the PCV field) in the PCV of 23h AP pupal wings. N = 8 (control) and 9 (MyoII-DN). \*\*\*\**P* < 0.0001. Data are mean + s.d. and were analysed by two-sided Mann-Whitney test. **f,** Adult wings in the PCV region in control (left, *en > mCD8::GFP*) and MyoII attenuated pupal wings (right, *en > MyoII-DN*). **g,** Effects of clonal expression of MyoII-DN within a subset of cells of the PCV field. pMad staining alone (white, left), or pMad staining (magenta), MyoII-DN expressing clones (green) at 25 h AP (right). Median filter applied to pMad staining. **h,** Top: Schematic depicting model of wild type pattern refinement whereby loss of cell shape changes from cells at the edge of the PCV field results in their exclusion from the field (by loss of BMP signalling and thus cell fate). Middle: Loss of MyoII activity and cell shape change throughout the PCV blocks refinement. Bottom: Loss of MyoII activity in a subset of PCV cells can lead to loss of signal and fate (ectopic refinement). **i,** Effects of clonal expression of Tkv^QD^ within a subset of cells of the PCV field. pMad staining alone (magenta, left), or pMad staining (magenta) Tkv^QD^-expressing clones (green) at 25 h AP (right). Median filter applied to pMad staining. **j,** Effects of Tkv^QD^ in *shi^ts1^* clones within a subset of cells of the PCV field. pMad staining alone (magenta, left), or pMad staining (magenta) *shi^ts1^* clones expressing Tkv^QD^ (green) at 24 h AP (right). Median filter applied to pMad staining. Scale bars: 50 μm for **b, d, f, i, j** and 25 μm for **g**.

What then is the role of cell shape changes in signalling pattern refinement? Despite reversal of BMP-induced cell shape changes being associated with reduced competence for vein fate, blocking cell shape changes did not affect BMP signalling. We hypothesised that what might be important is not cell shape change itself, but how cell morphology compares to that of other cells within the field. The impact of cell shape change loss may then be context-specific, facilitating refinement by causing less signalling and subsequent fate loss in cells surrounded by those with greater changes in shape. If this is the case, inhibition of cell shape changes in a small group of cells within the PCV field may decrease their ability to retain BMP signalling and vein fate. We tested this hypothesis by generating clones that attenuate cell shape changes amongst neighbours that are wild type.

Strikingly, when MyoII-attenuated clones are produced within the PCV field, loss of BMP signalling can often be observed in a context-dependent manner (Fig. 3g, Supplementary Fig. 2d, e). This suggests that MyoII-based cell shape changes play a crucial role in whether a cell retains vein fate during refinement, despite these shape changes not being required for BMP signalling. When all PCV field cells cannot form vein-like shapes, signalling still occurs throughout, and refinement does not take place (Fig. 3a-e, h). However, when cells are present in a heterogeneous population with or without cell shape changes, cells that can change shape both retain the signal and acquire vein fate (Fig. 3g, h). We propose that cells of the PCV field may be competing for BMP signalling via their changes in cell shape, with those that can change and retain vein-like shapes being more readily able to retain BMP signalling than their neighbours. Furthermore, our data indicate that the mechanism of pattern refinement is a self-organising mechano-chemical feedback loop, as BMP signalling induces the cell shape changes, which in turn influence the ability of cells to retain that signal.

### Competition for BMP signalling takes place at the PCV region

As BMP signalling appears to be needed for the refinement of the signal itself, we hypothesise that stronger signalling may make the cells more competitive for receiving the signal. Therefore, we generated strongly apically constricting clones expressing the constitutive active form of the BMP type I receptor (Tkv^QD^) and examined whether they perturbed endogenous BMP signalling at the PCV. Strikingly, wings carrying *tkv^QD^* clones often lose pMad signal in the PCV region, but not in LVs (Fig. 3i), suggesting that cells expressing Tkv^QD^ become super competitive for BMP signalling in a non-autonomous manner. This strongly supports the hypothesis that cells compete for BMP signalling in the PCV field, and that high levels of signalling in some cells are able to perturb signalling in others.

How do clones of ectopic BMP signalling non-cell-autonomously affect the ability of PCV region cells to receive the BMP signal? It is tempting to propose that they may capture extracellular Dpp ligand more readily, allowing them to out-compete others for the BMP signal. To address this, we generated clone cells expressing Tkv^QD^, but with less ability to capture the Dpp ligand. Since previous studies suggested that dynamin-mediated endocytosis plays a crucial role in BMP signalling and Dpp ligand capture in the wing imaginal disc^20^, we first tested whether this was also the case in the pupal wing. When clone cells of temperature-sensitive dynamin mutant (*shibire^ts1^; shi^ts1^*) are generated in the PCV region, pMad signal is often lost autonomously (Supplementary Fig. 3a), indicating that dynamin-mediated endocytosis is crucial for BMP signalling during PCV development. Next, we generated *shi^ts1^* clones co-expressing *tkv^QD^*. Although BMP signalling in these clones is still robust, BMP signalling in the PCV region is no longer disrupted (Fig. 3j). Taken together this indicates that cells with strong BMP signalling can outcompete other cells at the level of ligand capture through cell shape changes, decreasing their neighbours’ signalling capabilities.

### Basal cell shape changes facilitate competition between cells and pattern refinement

We next considered what the mechanism of competition between cells for the BMP signal is, and how cell shape might play a role. Previous studies indicate that extracellularly trafficked Dpp ligands are predominantly localised on the basal side of the wing epithelia^7^, therefore, the basal side of PCV field cells can be considered as the signalling micro-environment. We hypothesized that apical constriction may cause an expansion in basal cell size that could increase competence for capturing basally localised ligand. To investigate this, we compared the apical and basal cell sizes of cells both within and outside of the PCV field during the refinement period (Fig. 4a). We observe that as cells within the PCV field (BMP signal positive) apically constrict, they also basally expand, unlike cells outside of the PCV field (BMP signal negative) (Fig. 4b-d). Since the volume of cells is largely conserved throughout the refinement period (Supplemental Fig. 4a), it appears that basal expansion is induced by BMP-induced apical Myosin II without changing cell volume.

**Fig. 4:**
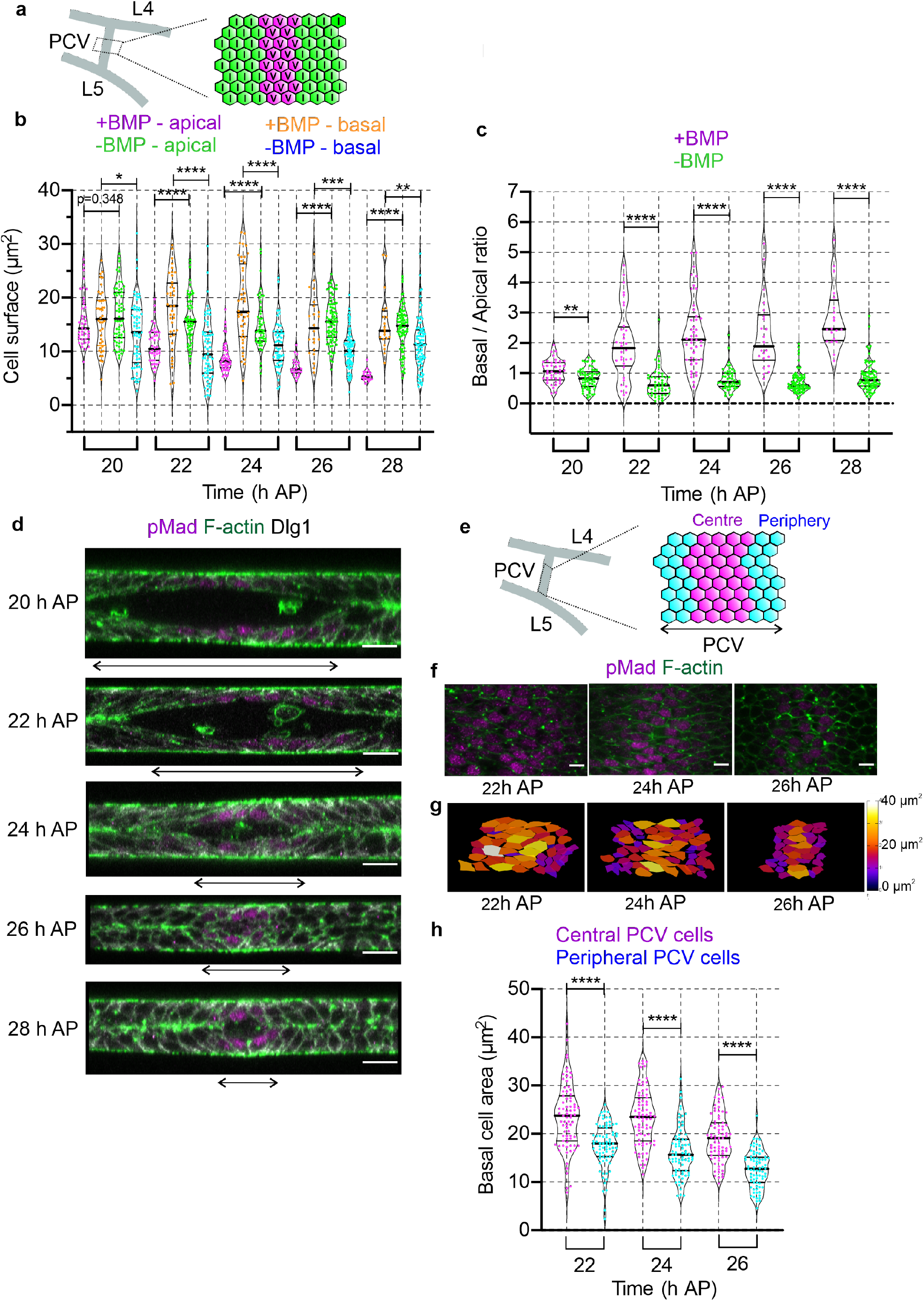
BMP signal induces apical constriction and basal expansion in the PCV field during PCV morphogenesis. **a,** Schematic of PCV field. Approximate position of cell area within and outside of the PCV field is shown as a square (left). Schematic of cells within (magenta) and outside of (green) PCV field during the refinement period. **b,** Apical and basal surface areas of cells within and outside of the PCV field during the refinement period. Sample sizes (from 5 wings) are 40 (20 h +BMP), 60 (20 h –BMP), 40 (22 h +BMP), 60 (22 h –BMP), 44 (24 h +BMP), 48 (24 h –BMP), 22 (26 h +BMP), 80 (24 h –BMP), 20 (28 h +BMP) and 79 (28 h –BMP). Violin plots show median, and 25th and 75th percentiles. Data from five independent images. Each data point [cells within the PCV: apical (magenta) and basal (orange); cells outside of the PCV: apical (green) and basal (cyan)] represents one cell. \**P* = 0.0329, \*\**P* = 0.0037, \*\*\**P* = 0.0002, \*\*\*\**P* < 0.0001. Data were analysed by two-sided Mann-Whitney test. **c,** Ratio of basal and apical surface of cells in **b**. Each data point [cells within the PCV (magenta) and cells outside of the PCV (green)] represents one cell. \*\**P* = 0.0030, \*\*\*\**P* < 0.0001. **d,** Optical cross sections focused on the PCV regions showing pMad (magenta) Dlg1 (white) and F-actin (green) at 20 h, 22 h, 24 h, 26 h and 28 h AP. Prospective PCV positions are indicated by double-headed arrows. **e,** Schematic of PCV field. Approximate position of cell area within the PCV field is shown as a square (left). Schematic of cells both central (magenta) and peripheral (cyan) PCV field during the refinement period. **f,** Max composites showing basal cell shape of PCV field cells at the level of central PCV cell basal surface during the refinement period. pMad staining (magenta) and F-actin (green) at 22h, 24h and 26h AP (left). Median filter applied to pMad staining. **g**, Heat map showing the changes in basal area of cells of the PCV field. The heatmap was produced using the ROI colour coder plugin, part of the BAR collection of ImageJ. **h,** Basal cell areas of peripheral and central PCV cells during refinement. N = 75 (15 cells per wing, data from 5 wings pooled). Violin plots show median, and 25th and 75th percentiles. Each data point (central PCV cells: magenta, peripheral PCV cells: cyan) represents one cell. \*\*\*\**P* < 0.0001. Data were analysed by two-sided Mann-Whitney test. The same wings are analysed for all of Figure 4. Scale bars: 10 μm for **d** and 5 μm for **f.**

We next investigated whether there was any difference in basal cell size between cells at the periphery of the PCV field, which may shortly lose vein fate competence, and the centre of the PCV field (Fig. 4e). We found that cells at the periphery often had smaller basal surfaces than those at the centre between 22 and 26 h AP, suggesting that cells which are to shortly lose their vein fate are less basally expanded (Fig. 4f-h).

To examine whether differential basal cell size is a mechanism by which cells could compete for the BMP signal, we observed whether differences in basal cell size within the PCV field are still observed when refinement does not take place due to attenuated MyoII activity. Our data reveal that, compared to wild type wings, there are much less pronounced differences in basal cell size between central and peripheral cells when MyoII is disrupted in the posterior half of pupal wing (Supplementary Fig 4b, c). These observations are consistent with basal cell size dynamics playing a role in the mechanism of refinement.

We next sought to ask how these changes in cell shape could make cells more competent for the BMP signal. A larger basal surface would allow a cell to occupy more space in the signalling microenvironment, however whether this would allow a cell to signal more would depend on how this impacted upon its ability to display the appropriate receptors. We investigated Tkv expression during PCV refinement, and found that Tkv localization is observed at the basal compartment in BMP positive cells, but not in negative cells (Fig. 5a, Supplementary Fig. 5a), suggesting that the cellular compartmentalisation of Tkv is regulated through a positive feedback mechanism.

**Fig. 5:**
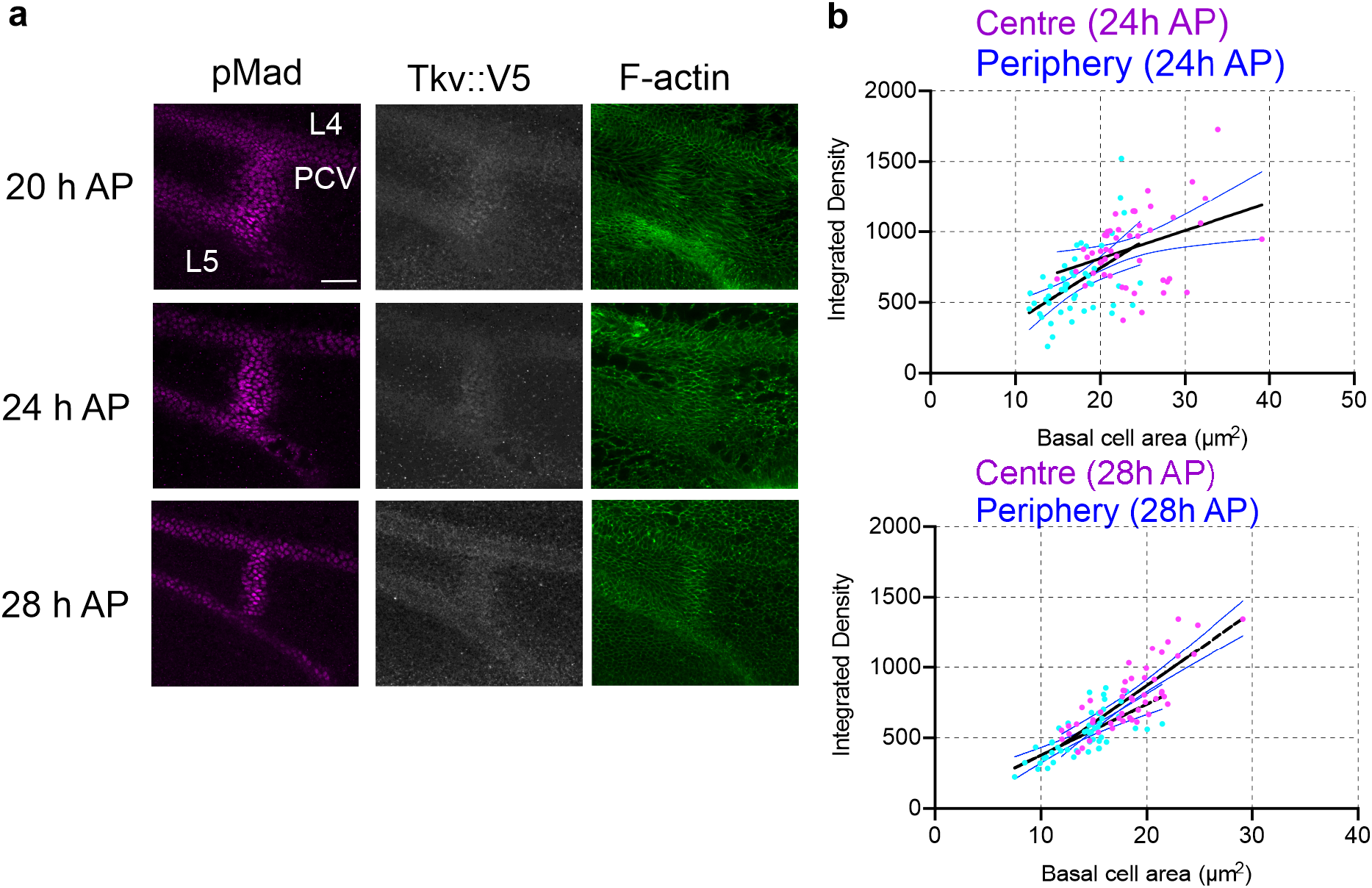
BMP receptor Tkv is basally localized in the PCV cells to sustain competence for BMP signalling. **a,** Tkv is localized at basal compartment in the PCV region during the refinement period. pMad expression (left, magenta), Tkv::V5 (middle, white) and F-actin (right, green) in the PCV region at 20 h, 24 h and 28 h AP. Median filter applied to pMad and, median and minimum filter applied to Tkv::V5. Images are maximum composites of basal region. Scale bar: 25 μm. **b,** Comparison of the integrated density of Tkv::V5 staining in the basal compartment and basal cell areas of peripheral and central PCV cells at 24 h and 28 h AP. Each data point (central PCV cells: magenta, peripheral PCV cells: cyan) represents one cell. N = 50 (10 cells per wing, data from 5 wings pooled). Black lines indicate mean levels and blue lines indicate error with 95% confidence intervals.

Correspondingly, Dpp ligands accumulate at the basal compartment in the PCV field during the refinement period (Supplementary Fig. 5b). Next, we quantified the interactions between basal cell area and integrated density of basally localized Tkv in the PCV field. Our data reveal that larger basal surface areas in PCV field cells correlated with higher overall intensities of Tkv, and that lower intensities of Tkv were observed for peripheral cells (Fig. 5b). Taken together, these results suggest that the mechanism by which vein-like cell shape changes facilitates cells to compete for the BMP signal is by promoting higher receptor localisation on the basal surface.

## Discussion

Here we found that cell shape changes are coupled to the refinement of BMP signalling during PCV morphogenesis. Our findings reveal that the formation of a mechano-chemical feedback loop drive competition for the vein fate-determining BMP ligand, with less competitive cells acquiring intervein cell fate. Therefore, different competence for the BMP signal leads to differential cell fate determination (vein or intervein) in later pupal development (Fig. 6a). Our data provide the evidence that clones of super competent cells are generated by expression of Tkv^QD^ (Fig. 3i) and that these can outcompete surrounding wild type cells and cause BMP signalling in the PCV field to be lost in a non-cell autonomous manner. The mechanism by which Tkv^QD^ clone cells dominate the BMP signal in the PCV field is not fully understood, however, Tkv^QD^ clone cells in *shi* mutants do not show such super competence (Fig. 3j), suggesting that high BMP signalling alone is not sufficient to increase competence. Converse results derived from context-specific expression of MyoII-DN provide further insights (Fig. 3). Basal cell shape changes via BMP signal-induced apical MyoII appear to be important for the determination of BMP positive and negative cells through generating differences among cells within the field, and may allow cells to compete for the extracellular ligand more efficiently. We propose that BMP signal-induced cell shape changes (via apical MyoII) provide the molecular mechanisms to increase competence for the BMP signal by forming a mechano-chemical feedback loop (Fig. 6b).

**Fig. 6:**
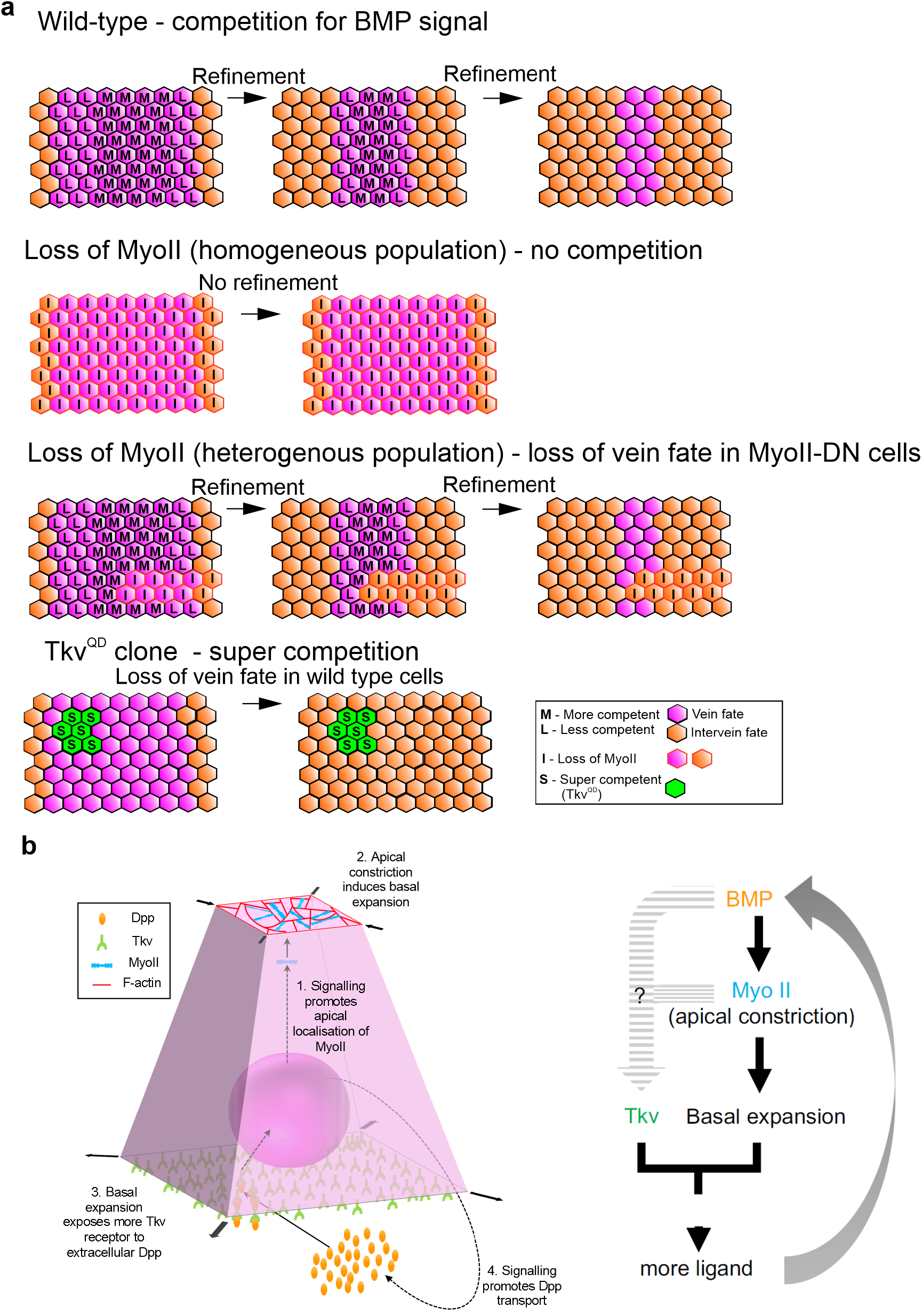
Schematic showing a mechano-chemical feedback loop in the PCV cells during refinement period. **a,** Schematics of competition for BMP signal during PCV development. Top: Schematic depicting model of wild type pattern refinement whereby loss of cell shape changes from cells at the edge of the PCV field results in the their exclusion from the field (by loss of BMP signalling and thus cell fate). Second row: Loss of MyoII activity and cell shape change throughout the PCV blocks refinement as no competition occurs. Third row: Loss of MyoII activity in a subset of PCV cells can lead to loss of signal and fate (ectopic refinement). Bottom: Clones expressing Tkv^QD^ lose BMP signalling in the PCV region in a non-autonomous manner, resulting in loss of PCV cell fate. **b,** Mechano-chemical feedback loop. 1. BMP signal promotes apical constriction through induction of apical Myosin II. 2. Apical Myosin II activity induces basal expansion. 3. Basal expansion exposes more basally localized Tkv receptors to extracellular Dpp. 4. Promoting BMP signal, feeding back into further shape change and promoting Dpp transport to the cell.

We propose that cells in the PCV field are in competition to receive the BMP signal during the refinement period. Our data suggest that basal surface dynamics form part of the mechanism of competition for the signal for the following reasons. First, the basal surface areas of PCV progenitor cells expand during the refinement period in a BMP dependent manner (Fig. 4). Second, basal surface sizes of peripheral PCV cells are significantly smaller than those at the centre of the PCV region (Fig. 4g, h). Third, Tkv sufficiently localizes at the basal compartment in the PCV region to capture basally trafficked Dpp ligand, and is present in greater quantities in basally expanded cells (Fig. 5, Supplementary Fig. 5). It is possible that cell shape changes play a role in promoting basal Tkv localisation, for example by downward cytoplasmic flow. Fourth, less significant differences of basal surface areas between the cells in the PCV region are noted when MyoII activity is disrupted (Supplementary Fig. 4b, c). Taken together, our observations indicate that BMP-induced apical MyoII leads to basal expansion, resulting into increasing competence for the BMP signal (Fig. 6). Previous studies have proposed that competition between cells for BMP signalling instructs the pattern of survival and elimination in the Drosophila wing imaginal disc^21–23^. Since BMP signalling is one of the key players regulating cell proliferation in the larval wing imaginal disc^24,25^, cells lacking BMP signalling are less proliferative than neighbouring cells and are eliminated as loser cells^26^. Although BMP still serves as a proliferative signal in the *Drosophila* wing during the early pupal stage^27,28^, BMP turns into a cell differentiation factor at the beginning of the refinement period and thus competition for it has a different outcome. However, unlike in classical cell competition, acquiring more Dpp ligand does not confer a selective advantage for survival, rather an alternative fate path (vein or intervein).

Unlike our recent observations that the 3D architecture of pupal wing epithelia (comprising of two apposing epithelial cell layers) and BMP signalling in the LVs are coupled^29^, PCV refinement appears to be a 2D phenomenon, as large single-layer clones expressing MyoII-DN that disrupt 3D architecture have been observed that do not affect refinement in the other layer (Supplementary Fig. 2e). This finding is intriguing, as we previously described that single-layer mutant clones of the Rho-GAP crossveinless-C (CV-C), which promotes Dpp ligand transport, could affect BMP signalling in the PCV region of the other layer^6^. These phenomena are likely to be distinct, as unlike when expressing MyoII-DN, small clones of *cv-c* mutants do not lose BMP signalling, whereas larger ones do. These results suggest that the inter-layer effects previously observed for *cv-c* mutant clones require the proper apposition of the two wing layers, which is locally disrupted in both MyoII-DN clones and tissue-level MyoII-DN expression (Supplementary Fig. 4d).

How cell shape-based competition for Dpp ligand allows refinement to form a neat stripe of vein cells is still not clear. We postulate that cells surrounded by other BMP-positive apically constricted cells will be more readily able to form a vein-like shape themselves, a phenomenon likely to be amplified by the mechanical feed-forward loop we have identified. This could explain why cells at the edge of the field, next to intervein cells, are less able to expand basally, and thus are more likely to lose BMP signalling competence, leaving a stripe of vein-fated cells. Non-cell autonomous feed-forward factors downstream of BMP signalling, such as Cv-C, could also play a role, as optimising the extracellular transport of Dpp affects not only an individual cell but also other cells nearby. It is tempting to speculate whether differences in signalling induced by differential cell shape changes can play a role in the refinement of the extracellular ligand range, reinforcing differences in cell shape. Notably, not all peripheral cells are smaller than all central cells at any one snapshot (Fig. 4g). This is consistent with the PCV field not refining uniformly, with different sides or positions within the field refining at different times (Fig. 2c). However, it could also be the case that smaller central cells are able to persist because of an accommodating environment generated by basally expanded cells nearby.

We hypothesize that cell shape-based competition for an extracellular signalling component is likely to be a general mechanism for self-organisation of pattern refinement during development. Cell shape changes are a common part of the morphogenesis programme and could feed back into developmental patterning in a variety of contexts^3,4^. Apical constriction and basal expansion are an important aspect of epithelial folding, a process which has broadly been linked to cell fate decisions and developmental patterning^3,4,30,31^. Our finding that cell shape changes within the 2D epithelial layer, irrespective of epithelial folding, can instruct pattern refinement provides a novel insight into epithelial morphogenesis.

In summary, our data reveal that cell shape changes influence refinement of the signalling pattern by facilitating cells to compete for signalling pathway activation. We have uncovered that competition for extracellular BMP ligand occurs via a mechano-chemical feedback loop between cell shape changes and BMP signalling, leading to self-organising refinement of the developmental field during pattern formation.

## Acknowledgements

We are grateful to Jukka Jernvall, Charlotte Repton and Yukitaka Ishimoto for thoughtful comments on the manuscript. We thank the Light Microscopy Unit of the Institute of Biotechnology of the University of Helsinki for their support. We thank G. Pyrowolakis, H. Ohkura, R. Le Borgne, D. Kiehart, T. Kornberg and A. Martin for fly stocks. This work was supported by grant 308045 from the Academy of Finland, the Sigrid Juselius Foundation to O.S., grant 295013 from the Academy of Finland to D.T-M., and the Center of Excellence in Experimental and Computational Developmental Biology from the Academy of Finland to O.S. and I.S-C.

## Author contributions

D.T-M. and O.S. conceived the project and planned experiments. D.T-M., M.P.M., N.V.T. and H.A. performed experiments. D.T-M., M.P.M., N.V.T., E.M.B. and H.A. analysed the results and discussed them with O.S. I.S-C. provided inputs. D.T-M. and O.S. wrote the manuscript and all authors made comments. O.S. supervised the project.

## Data availability

The datasets generated and analysed during the current study are available from the corresponding authors on reasonable request.

## Materials and methods

### Fly genetics

*UAS-mCD8::GFP* (#5137), *en-Gal4* (#30564), *shi^ts1^* (#7068) and UAS-MBS.N300 (#63791) were obtained from the Bloomington *Drosophila* Stock Centre. *UAS-tkv^Q253D^* and *cv^70^* were described previously^8,10^. *E-cad::GFP* was obtained from H. Ohkura^17^, *MyoII::RFP* from R. Le Borgne^15^, *UAS-MyoII-DN* from D. Kiehart^18^, *Tkv::V5* from G. Pyrowolakis^32^, *Dpp::mCherry* from T. Kornberg^33^ and *sqh-GAP43::mCherry* from A. Martin^34^. Populations of mixed sex were used except for when using *yw* or *tubP-Gal80^ts^* on X, where females were selected, and experiments involving *crossveinless*, where only males were used. The ages of pupal wings at dissection are given at developmental timepoints equivalent to 25°C. Calculations for relative developmental timing at 18°C, 25°C and 29°C were based on previously published data and rounded to the nearest hour^35^. For experiments using *en-Gal4*, pupae were raised at 18°C for 22 hours after pupariation, and then shifted to 29°C for either 10 (Fig. 3b, c, d, e, Supplementary Fig. 2a, 4d) or 12 (Supplementary Fig. 4b-c) hours before dissection and fixation. The exception to this was the experiment involving UAS-MBS.N300, when pupae were raised at 29°C for 21 hours after pupariation (Supplementary Fig. 2b). For clone generation, larvae were raised at 25°C for 3-4 days AEL (or 18°C for 6-7 days AEL), before being heat shocked in a 37°C water bath for 1 hour. Vials containing larvae were then either placed at 18°C until at the white pre-pupal stage, and then transferred to 29°C for 21 hours before dissection and fixation (Fig. 3g, i, Supplementary Figs 1b, 2d, e) or transferred to 25°C for 20 hours after pupariation and then 34°C for 3hours (Fig. 3j, Supplementary Fig. 3). For time-lapse imaging, pupae were raised at 25°C until 17 hours after the pre-pupal stage. They were then moved to room temperature for one hour, during which windows were cut into the pupal case and pupae mounted, before being imaged as previously described^29^.

### Full genotypes

Fig. 1d-e, Fig. 4 and Supplementary Fig. 4a : *yw*

Fig. 1f: *yw; ubi-E-cad::GFP,* or *cv^70^; ubi-E-cad::GFP*

Fig. 1g and Supplementary Fig. 1a: *MyoII::RFP,* or *MyoII::RFP, cv^70^*

Fig. 2 and Supplementary videos 1-3: *ubi-E-cad::GFP/sqh-Gap43::mCherry*

Fig. 3e-f and Supplementary Fig. 4d: *en-Gal4/UAS-mCD8::GFP; tubP-Gal80^ts^,* or *en-Gal4/UAS-MyoII-DN; tubP-Gal80^ts^*

Fig. 3g and Supplementary Fig. 2d, e: *hs-Flp; tubP-Gal4 UAS-mCD8::GFP/UAS-MyoII-DN; tubP-Gal80 FRT^82B^ /FRT^82B^*

Fig 3i: *hs-Flp FRT^82B^; tubP-Gal4 UAS-mCD8::GFP/UAS-tkv^Q253D^; FRT^82B^ tubP-Gal80/FRT^82B^*

Fig 3j: *hs-Flp, tubP-Gal80, FRT^19A^/shi^ts1^, FRT^19A^; tubP-Gal4, UAS-mCD8::GFP/UAS-tkv^Q253D^*

Fig 5 and Supplementary Fig 5a: *yw; tkv-V5/CyO*

Supplementary Fig. 1b: *hs-Flp/tubP-Gal80^ts^, MyoII::RFP; tubP-Gal4 UAS-mCD8::GFP/UAS-tkv^Q253D^; tubP-Gal80 FRT^82B^ /FRT^82B^*

Supplementary Fig. 2a: *cv^70^; en-Gal4/+; tubP-Gal80^ts^*, or *cv^70^; en-Gal4/UAS-MyoII-DN; tubP-Gal80^ts^*

Supplementary Fig. 2b, c: *en-Gal4/+; UAS-mbs.N300/tubP-Gal80^ts^ or en-Gal4/UAS-mCD8::GFP; tubP-Gal80^ts^*

Supplementary Fig. 3: *hs-Flp, tubP-Gal80, FRT^19A^/Shi^ts1^, FRT^19A^; TubP-Gal4, UAS-mCD8::GFP*

Supplementary Fig. 4b-c: *en-Gal4/sqh-Gap43::mCherry; tubP-Gal80^ts^*, or *en-Gal4/UAS-MyoII-DN, sqh-Gap43::mCherry; tubP-Gal80^ts^*

Supplementary Fig. 5b: *dpp::mCherry* (II)

### Immunohistochemistry

Pupae were fixed in 3.7% formaldehyde (Sigma-Aldrich) for two nights at 4°C before dissecting the pupal wings and blocking with Normal Goat Serum (10%) overnight. Both primary and secondary antibody incubations also took place overnight at 4°C. The following primary antibodies were used: mouse anti-DLG1 [1:40; Developmental Studies Hybridoma Bank (DSHB), University of Iowa] and rabbit anti-phospho-SMAD1/5 (1:200; Cell Signaling Technologies) mouse anti-V5 (1:200, Invitrogen). Secondary antibodies were anti-rabbit IgG Alexa 568 (1:200, Invitrogen), anti-mouse IgG Alexa 647 (1:200; Life technologies), anti-rabbit IgG Alexa 647 (1:200; Life technologies). F-actin was stained with Alexa 488 conjugated phalloidin (1:200; Life technologies).

### Imaging and Image Analysis

Confocal images and time-lapse images were acquired using a Leica SP8 STED confocal microscope. Time-lapse images were processed using Imaris v9.1.2 (Bitplane/Oxford Instruments) and snapshots segmented by hand in Image J/Fiji (v. 1.52p and 1.53c)^36^. All other images were processed and analysed using Fiji. All images (with the exception of cross sections), including time-lapse snap shots, are maximum composites. A median filter was applied to pMad images, and median and minimum filters were applied to Tkv::V5 images in Fiji. The heatmaps were generated using the ROI color coder plugin, part of the BAR collection of ImageJ^37^.

The number of pMad-positive cells within the PCV field was calculated by first excluding adjacent LV nuclei by marking the predicted trajectory of the LV–PCV boundary by drawing across from the edge of L4 and L5 on either side of the PCV. pMad positive nuclei between these lines were then counted. Z projections of median filtered images were used for quantification, using the stacks for reference.

The apical and basal sizes of field cells were analysed using individual slices of stacks in Fiji. Cells that showed pMad staining in the nucleus were traced and measured. The most apical or basal slice where the cell outline was clear was used for each cell. Cells at the centre and posterior portion of the PCV were analysed. Cells in the middle 50% of the PCV (as indicated by pMad) were designated central cells and those outside this peripheral.

Cell volumes were determined for the same cells for which apical and basal surface areas had been measured. To measure cell volumes, the RoiManager3D tool was used, a plug-in in the 3D ImageJ Suite (version 3.96)^38^. In any z-sections for which cell boundaries were ambiguous, the boundaries were drawn as best-guess interpolations between the segmented boundaries of neighboring apical and basal z-sections. Cells with more than one consecutive ambiguous z-section boundary were excluded from the subsequent volume determinations.

Figure 1g and Supplementary Fig 1a were generated from the same stacks. Figure 4 and Supplementary Figure 4a represent analyses of the same data set, as do Figure 3b-e and Supplementary Figure 4d, and Figure 5 and Supplementary Figure 5a, respectively. All representative images are representative of at least three biological replicates.

### Statistics

Statistical analyses were performed using GraphPad Prism software (v.8.3.0, GraphPad). The number for all quantified data is indicated in the figure legends. All *P* values were calculated using a two-sided Mann-Whitney test and specified in the figure legends and in the corresponding plots.

