## Supplemental figures for "Mechano-chemical feedback leads to competition for BMP signalling during pattern formation"

**Supplementary figures**

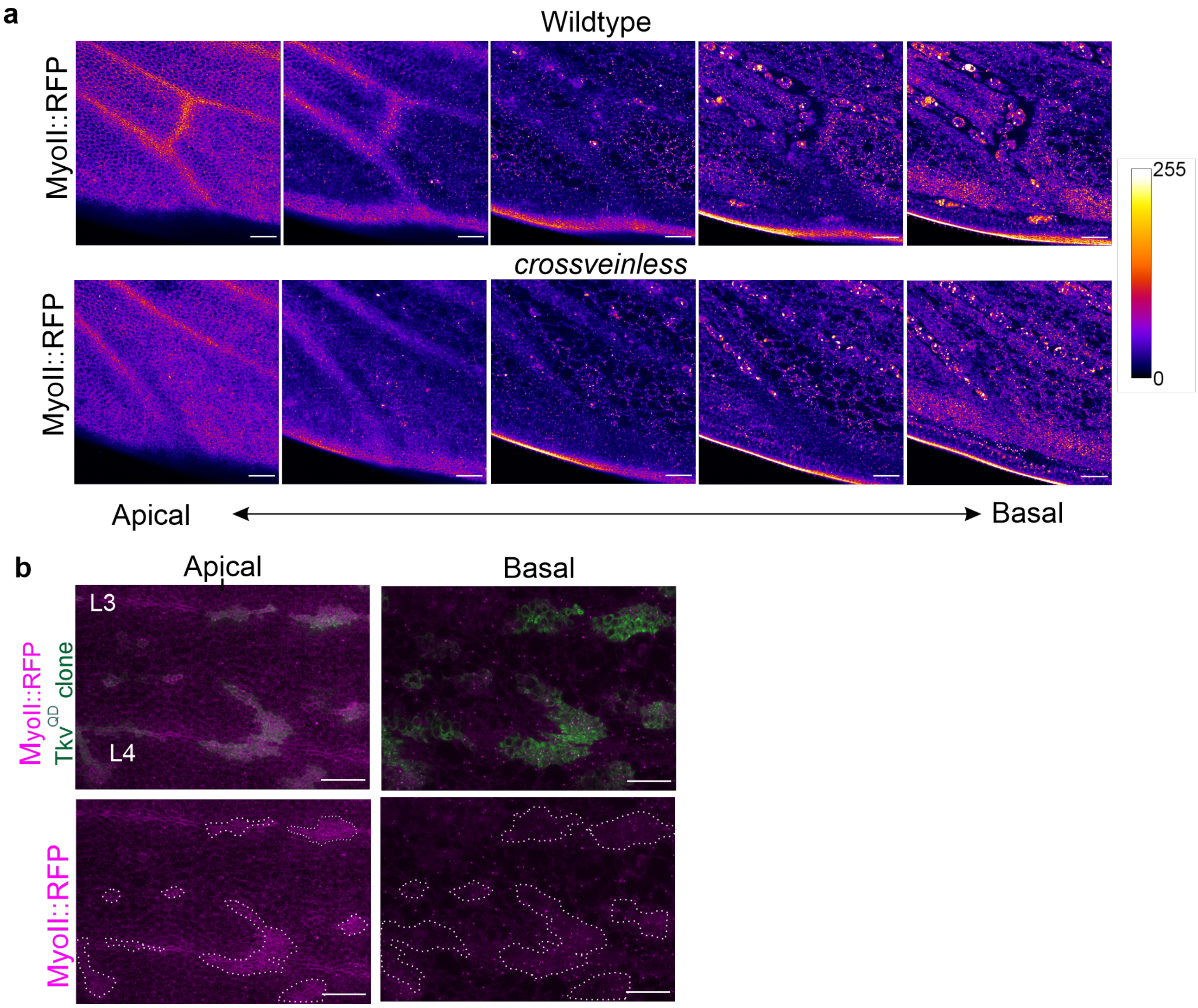


**Supplementary Fig. 1: BMP signal induces apical enrichment of Myosin II in pupal wings. a,** MyoII-RFP localisation throughout progressive apical to basal projections of PCV field and intervein cells in wild type and *crossveinless* mutant 24 h AP pupal wings. Maximum composites of five different sequential sections of the pupal wing along the apicobasal axis. Same tissues shown as in Fig 1g. **b,** Apical and basal maximum composites showing MyoII-RFP localisation (magenta) in ectopic clones expressing constitutively active BMP receptor $Tkv^{QD}$ (green) (25 h AP). Dashed lines depict overexpression clones. Scale bars: 25 µm for **a** and **b**.

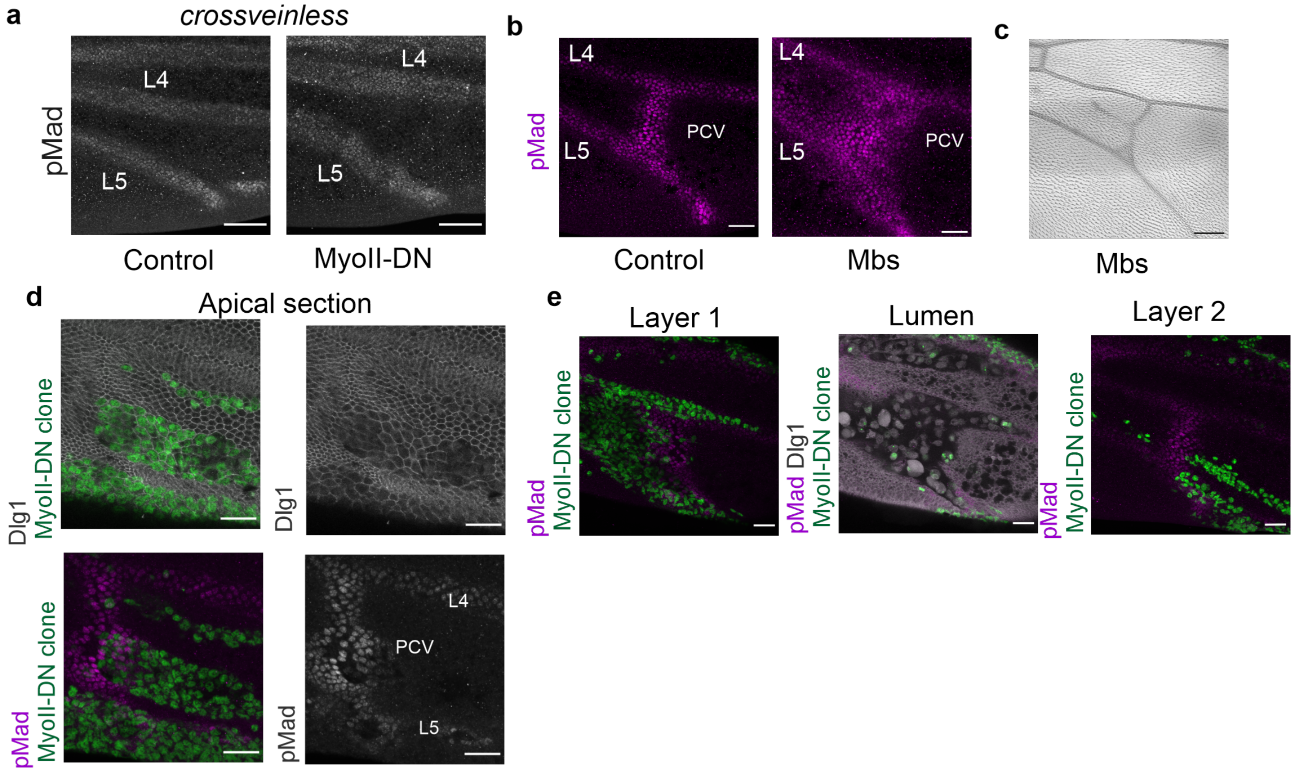


**Supplementary Fig. 2: MyoII activity affects BMP signalling pattern refinement in the PCV field. a,** pMad expression in the PCV region in control (left, *en-Gal4*) and MyoII attenuated pupal wings (right, *en > MyoII-DN*) at 23 h AP in a *crossveinless* background. Median filter applied. **b,** pMad expression in the PCV region in control (left, *en > mCD8::GFP*) and Mbs overexpression (right, *en > Mbs*) at 25 h AP. Median filter applied. **c,** Adult wing in the PCV region in Mbs overexpressing pupal wing (right, *en > Mbs*). **d,** Effects of clonal expression of MyoII-DN within a subset of cells of the PCV field on pMad staining (magenta - bottom) and apical cell size (Dlg1 – top) at 25 h AP. MyoII-DN expressing clones are marked by GFP (green). Note that large clones contain cells that both lose and retain the BMP signal despite attenuation of MyoII activity. Median filter applied to pMad staining. **e,** Effects of large clones expressing MyoII-DN (left) that disrupt apposition of cell layers and lumen formation (middle), on the other cell layer (right) at 25 h AP. Median filter applied to pMad staining. Scale bars: 50 μm for **a, c**, 25 μm for **b, d, e**.

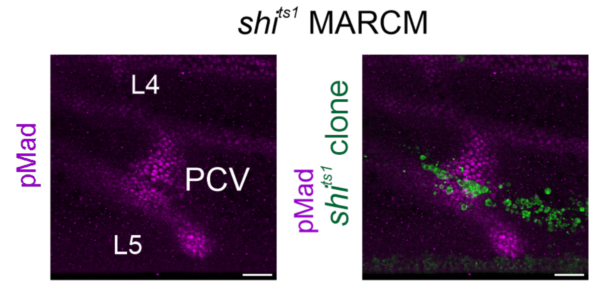


**Supplementary Fig. 3: Dynamin-mediated endocytosis is crucial for BMP signalling in the PCV field.** Effects of *shi*$^{ts1}$ clones within a subset of cells of the PCV field. pMad staining alone (magenta, left), or pMad staining (magenta) *shi*$^{ts1}$ clones (green) at 24 h AP. Median filter applied. Scale bars: 25 µm.

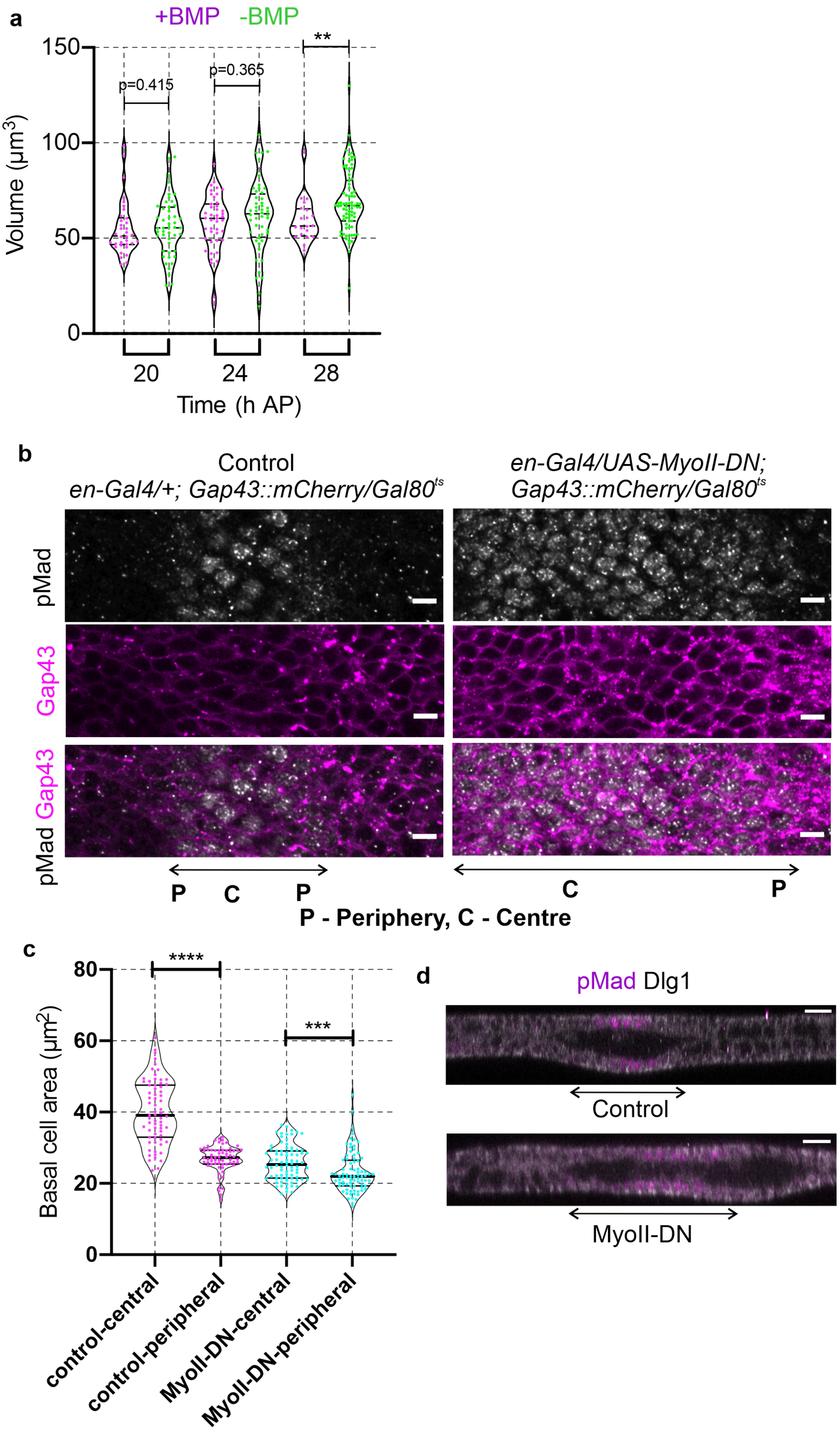
a
+BMP -BMP
Volume (µm3)
150
100
50
0
p=0.415
p=0.365
**
20
24
28
Time (h AP)
b
Control
en-Gal4/+; Gap43::mCherry/Gal80ts
en-Gal4/UAS-MyoII-DN;
Gap43::mCherry/Gal80ts
pMad
Gap43
pMad Gap43
P C P
C P
P - Periphery, C - Centre
c
Basal cell area (µm2)
80
60
40
20
0
****
***
control-central
control-peripheral
MyoII-DN-central
MyoII-DN-peripheral
d
pMad Dlg1
Control
MyoII-DN

31

**Supplementary Fig. 4 Loss of MyoII activity attenuates BMP-induced basal expansion in the PCV field during PCV morphogenesis: a,** Volume of cells within and outside of the PCV field during the refinement period. Sample sizes (from 5 wings) are 34 (20 h +BMP), 47 (20 h –BMP), 43 (24 h +BMP), 48 (24 h –BMP), 20 (28 h +BMP) and 79 (28 h –BMP). Violin plots show median, and 25th and 75th percentiles. Each data point [cells within the PCV (magenta) and cells outside of the PCV (green)] represents one cell. **$P = 0.0047$. Data were analysed by two-sided Mann-Whitney test. Wings are from the same dataset as Fig. 4. **b,** Basal cell shapes (Gap43 - magenta) in the PCV field (marked by pMad – white) in control (left) and MyoII attenuated pupal wings (right, *en* > *MyoII-DN*) at 25 AP. Maximum composite of basal regions of the cells are shown. Prospective PCV positions are indicated by double-headed arrows, as well as the positions of the middle (M) and periphery (P) of the PCV field. Median filter applied to pMad staining. **c,** Basal cell areas of peripheral and central PCV cells in control and MyoII attenuated pupal wings (*en* > *MyoII-DN*) at 25 h AP. N = 75 (15 cells per wing, data from 5 wings pooled). Violin pots show median, and 25th and 75th percentiles. Each data point (central PCV cells: magenta, peripheral PCV cells: cyan) represents one cell. ***$P = 0.0004$, ****$P < 0.0001$. Data were analysed by two-sided Mann-Whitney test. **d,** Optical cross section focused on the PCV region of showing control (top, *en* > *mCD8::GFP*) and MyoII attenuated pupal wings (bottom, *en* > *MyoII-DN*) at 23 h AP. BMP signal positive positions are indicated by double-headed arrows. Scale bars: 5 µm for **b** and 20 µm for **d**.

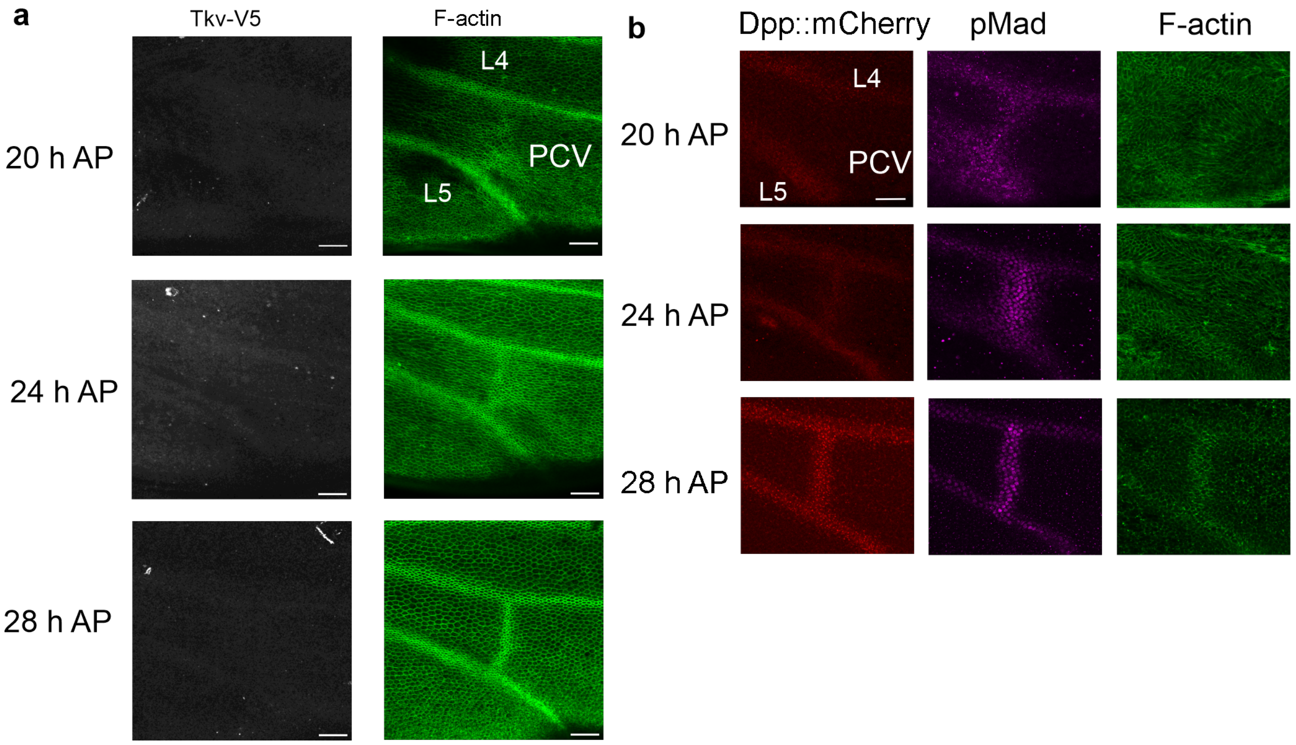


**Supplementary Fig. 5 BMP signalling components localise in the basal compartment: a.** Apical compartment of Tkv::V5 (left, white) and F-actin (right, green) in the PCV region at 20 h, 24 h and 28 h AP. Images are maximum composites of apical regions. Median and minimum filter applied to Tkv::V5. Scale bar: 25 µm. **b.** Dpp accumulates at basal compartment in the PCV region during refinement period. Dpp::mCherry (left, red), pMad (middle, magenta) and F-actin (right, green) in the PCV region at 20 h, 24 h and 28 h AP. Median filter applied to pMad. Images are maximum composites of basal region. Scale bar: 25 µm.

**Supplementary videos**

**Supplementary Video 1:** Time-lapse imaging of E-cad::GFP in the PCV region between 18.5 h and 29.5. h AP. Three clusters of cells are marked. Future PCV cells (magenta) show progressive apical constriction. Cells immediately adjacent to future PCV cells (cyan) show transient apical constriction during 22 h and 26 h AP and revert to intervein-like conformation. Cells distant from PCV cells do not show apical constriction through the time-lapse imaging. The same cells are shown in supplementary video 1 and 2, and Figure 2a and c.

**Supplementary Video 2:** Time-lapse imaging of E-cad::GFP in the PCV region between 24 h and 28 h AP. Cells marked by magenta and cyan show apical constriction at 24h AP. In contrast, cells only marked by magenta maintain apical constriction at 28h AP, suggesting that cells marked by cyan fail to maintain vein-like morphogenesis. The same cells are shown in supplementary video 1 and 3, and Figure 2a and c.

**Supplementary Video 3:** Time-lapse imaging of E-cad::GFP in the PCV region between 19h, and 21 h AP prior to apical constriction of future vein cells. Please note that future PCV (magenta) and intervein cells (cyan) are within the same cell lineage at the early PCV morphogenesis. The same cells are shown in supplementary video 2 and 3, and Figure 2a and c.
